# Purging of inbreeding depression does not eliminate environmental variation in reproductive onset

**DOI:** 10.64898/2026.03.11.711066

**Authors:** Sarthak Grover, Elise Jeanne, Steven A. Ramm

## Abstract

Many simultaneous hermaphrodites use selfing for reproductive assurance only when outcrossing opportunities are limited, owing to inbreeding depression in selfed progeny. However, scenarios that enforce substantial selfing (such as during recolonisation) can rapidly select for a high selfing propensity, a shift in mating system that is expected to eliminate both inbreeding depression and the delayed reproductive onset under selfing that is typically associated with it. We tested these predictions in the flatworm *Macrostomum hystrix*, using a line derived from an outcrossing population that had been subjected to enforced selfing for multiple generations followed by several years of relaxed selection. As predicted, isolated individual forced to self and individuals with constant partner access (i.e. outcrossing opportunities) did not differ in reproductive onset nor in inbreeding depression estimated through offspring survival. However, a third treatment group that provided intermittent partner access (to allow outcrossing but minimise potential competition effects) showed a different pattern: no inbreeding depression in offspring but a substantially accelerated reproductive onset. Whilst our results thus support the effective purging of inbreeding depression and increased selfing propensity under enforced selfing, we suggest that cues of an unstable social or physical environment nevertheless exert a major influence on reproductive timing.

## Introduction

Self-fertilisation (selfing) can provide reproductive assurance when outcrossing opportunities are scarce [1-4], but is typically associated with inbreeding depression [5]. To balance these fitness benefits and costs, many organisms have therefore evolved facultative strategies that regulate selfing rates, as for example in the diverse range of taxa that exhibit delayed selfing behaviour [6-10]. In this strategy, individuals prefer to outcross, but switch to selfing in the absence of outcrossing opportunities only after a delay period (called the ‘waiting time’) has passed [11]. This added delay allows individuals to (1) increase their chances of finding an outcrossing opportunity and thereby (2) avoid wasting resources on less fit inbred progeny if an outcrossing opportunity arises between reproductive maturity and the end of the waiting period.

Selfing results in inbreeding depression because it increases homozygosity [12]. In zygotes formed by selfing, both male and female gametes originate from the same parent individual, and this increases the chances of passing on two copies of the same allele at each locus, including recessive deleterious alleles that then lower fitness when expressed in selfed progeny [5]. Theory therefore predicts that the optimal duration of the waiting time will scale with the magnitude of inbreeding depression: the higher the costs of selfing, the more reluctant the individual should be to self [11].

Inbreeding depression costs strongly depend on a population’s history of inbreeding. In preferentially outcrossing populations, where selfing occurs occasionally for reproductive assurance, recessive deleterious alleles are rarely expressed and exposed to selection and therefore remain segregating in the population. Consequently, every selfing event results in high fitness loss and regular selfing is normally expected to be selected against [13, 14].

However, certain ecological conditions — such as range expansions [15, 16] or re-colonisation following population fragmentation [17] or bottlenecks [18] — can put a high premium on reproductive assurance, resulting in successive generations of selfing. Such regular selfing has an important evolutionary consequence — it reduces inbreeding depression costs over time. When populations self regularly, segregating recessive deleterious alleles are either purged by selection [19] or fixed by drift [20], progressively lowering inbreeding depression costs over successive generations. As these costs decrease, selection should favour shorter waiting times [11, 21], indicating an increased selfing propensity. While many studies support an association between inbreeding depression magnitude and waiting time duration [6, 8, 9, 22-24] or between realised selfing rates and observed inbreeding depression [6, 25, 26] (presumably reflecting historical purging), to our knowledge only one so far has experimentally demonstrated that regular selfing lowers inbreeding depression costs and consequently reduces waiting times under controlled laboratory conditions, in the freshwater snail *Physa acuta* [19].

Here, we take advantage of a highly inbred line of a free-living flatworm, produced by repeated generations of self-fertilisation, to test whether regular selfing indeed lowers inbreeding depression costs and increases selfing propensity. Specifically, we measured hatchling-to-adult survival and waiting times in a *Macrostomum hystrix* line that was derived from a preferentially outcrossing population with significant inbreeding depression [8], but then produced by 8 generations of selfing and multiple generations of matings between close relatives (expected F > 0.998 in 2011), followed by c. 15 years (>100 generations) of being maintained at larger population sizes (often >1000 individuals), i.e. under relaxed selection [27]. After such a strong period of inbreeding, we thus expect to find no inbreeding depression and no waiting times in this line, even though the ancestral outcrossing population exhibited both [8], unless either a) the purging process was incomplete, meaning residual recessive deleterious alleles could still cause inbreeding depression and waiting time in this line; or b), inbreeding depression has been re-generated due to deleterious recessive mutations accumulated in the intervening years, both of which we considered unlikely.

Waiting times and inbreeding depression in *Macrostomum* species have thus far been measured by comparing the age at reproductive onset and progeny fitness in two treatment groups: one where individuals are isolated and forced to self, and the other where mating partners are always grouped, ensuring constant outcrossing opportunities [8, 22, 28]. In addition to these treatments, we included in this experiment an additional ‘intermittent grouping’ outcrossing treatment that provided regular outcrossing opportunities while substantially limiting the time the mating partners were grouped together. We could thereby distinguish the effects of outcrossing opportunities from extended grouping *per se*. Indeed, constant access to mating partners in experimental setups might not reflect the ecological reality for certain species and could even be stressful, with at least two previously reported effects for hermaphroditic animals likely to be relevant here. First, in certain freshwater snails, constant grouping is thought to create resource competition that reduces fitness [29].

And second, in the sea slug *Alderia willowi*, a similarly reduced fitness directly results from the inflated mating rates that occur under constant grouping [10]. *A. willowi* sperm are transferred by traumatic insemination, so each insemination event likely imposes an energetic cost to the recipient in terms of tissue repair. An important second goal of our experiment was therefore to compare treatments with constant versus periodic outcrossing opportunities, to test whether any of these effects exist for *M. hystrix*, which also reproduces by traumatic hypodermic insemination, injecting sperm directly into the body wall of conspecifics [30] or itself [31] during outcrossing and selfing events, respectively.

## Methods

### Study organism

*Macrostomum hystrix* Ørsted 1843 sensu Luther 1905 is a widely distributed free-living, hermaphroditic flatworm found in brackish habitats, with previously studied populations in the Baltic and the Mediterranean seas [8, 28, 30, 32, 33].

Initially determined to be a preferentially outcrossing species with delayed selfing [8], *M. hystrix* populations in fact show wide natural variation in their mating system expression, including in waiting times. Isolated individuals from ‘BIB’ and ‘TVÄ’ populations (collected in Bibione, Italy and Tvärminne, Finland, respectively) self at the same time as grouped individuals [28], but those in the ‘PISA’ population (collected from the San Rossore Regional Park near Pisa, Italy) switch to selfing only after a substantial delay – 54%, on average, though there was substantial genetic variation for selfing propensity [8], in line with previous estimates in *P. acuta* [24, 34].

This study used the highly inbred SR1 line, which was derived from a single founding individual from the ‘PISA’ population that showed the earliest selfing behaviour followed by multiple generations of selfing and sib-sib mating, meaning the inbreeding coefficient (*F*) of this line reached an estimated 0.998 by 2011. After such strong inbreeding, the deleterious alleles responsible for causing inbreeding depression are expected to be either purged or fixed, effectively eliminating any fitness differences between selfed and outcrossed offspring (i.e. inbreeding depression). Since then, the SR1 line has been regularly bottlenecked and maintained under standard lab conditions (20°C, 6‰ or 12‰ artificial sea water (ASW), 14:10 light:dark cycle) under relaxed selection for selfing propensity.

### Generation of experimental individuals

We left c. 100 adults to lay eggs in Petri dishes for 5 days and harvested hatchlings daily. These experimental hatchlings were allocated to their respective treatments by transferring them to wells of 24-well cell culture plates (Geyer LABSOLUTE 24-well Zellkulturplatten Art. 7 696 792) containing 1.5mL of 12 ‰ ASW per well. Worms were fed *ad libitum* on *Nitzschia curvilineata* diatoms throughout the experiment. At each feeding, age-matched worms received 50 µL of a dense algal suspension per individual.

### Treatments

To simulate varying opportunities for outcrossing versus selfing, we implemented three experimental treatments: Isolated, Triplet and Intermittently Grouped. We started with 32 replicates per treatment that were distributed in a balanced fashion across eight 24-well plates. Each Isolated (I) replicate consisted of one individual per well and all I individuals were thus forced to self in order to reproduce. Each Triplet (T) replicate consisted of three individuals per well, always grouped together and thus with constant outcrossing opportunities. By contrast, each Intermittently Grouped (IG) replicate consisted of three individuals housed separately in individual wells for most of the experimental period, except for regular grouping periods (2 hours every 2^nd^ or 3^rd^ day) in a fourth ‘neutral’ well that provided an opportunity for outcrossing with two conspecifics assigned to the same replicate. During each grouping event, we transferred all three worms to the designated neutral well for 2 hours, then returned them to individual housing wells. There was no way of knowing which housing well each individual worm came from, so in practice housing wells were shuffled following every grouping session. To minimise handling differences between treatments, I and T individuals were also pulled into the micropipette tip (used to manipulate worms) once at the beginning of grouping and once at the end, but returned to their same well after a minimum pause of 1 second. Grouping for IG replicates started from the second day after hatchlings were allocated to treatments and continued until the experiment ended. Individuals were grouped every 2^nd^ day until they were 64-69 days old, after which the interval was adjusted to every 3^rd^ day for logistical reasons.

Note that it was not possible to determine the proportion of individuals that actually outcrossed in either outcrossing regime, as paternity assays cannot be conducted on the highly inbred SR1 line due to the expected lack of genetic variation in microsatellite markers.

However, individuals in this line still retain a male-biased sex allocation (the relative investment into male versus female function) and the ability to plastically proportion more resources to the male function in the presence of mating partners [27]. These traits are typically found in outcrossing *Macrostomum* species [35]. In contrast, the congener *Macrostomum pusillum*, estimated to be an regularly selfing species, shows a weaker allocation to male function and lacks the ability to plastically adjust its sex allocation [22]. Both high investment into the male function and the ability to plastically increase it in response to group size are likely wasteful if the individual is ultimately selfing.

### Observations

We recorded age at reproductive onset by monitoring all wells for the presence of hatchlings. Wells were monitored daily from the start of the experiment until day 72, thereafter every second day until day 93, and then every third day from day 94 until the end of the experiment on day 137. Note that for logistical reasons, no groupings or observations were performed from day 76 to day 82 of the experiment. In order to monitor offspring survival until adulthood, we collected up to 5 hatchlings for each replicate on the first day they were observed and housed them in individual wells of 48-well plates. Survival of these hatchlings was checked when they would have been at least 21 days old. Hatchlings not found in their wells or found undeveloped on check were also considered as dead (i.e. failed to achieve sexual maturity) for the purposes of the analysis.

### Statistical analyses

All statistical analyses were performed, and figures were created, using R version 4.2.2 [36] and RStudio [37]. To assess the effects of treatment on age at reproductive onset, a linear mixed effects model (LMM) was fitted using the ‘lme4’ package with each replicate’s plate ID included as a random effect. A Type III ANOVA was applied to the model using the ‘car’ package to test for significance of the fixed effects and post-hoc pairwise comparisons were performed using Tukey-adjusted pairwise comparisons via the ‘emmeans’ package. Normality and homoscedasticity assumptions were assessed using residual plots.

To assess whether the survival of collected hatchlings differed by treatment, we first calculated the proportion of hatchlings that survived per replicate as the dependant variable and then fitted a linear model using the lm() function with treatment as the fixed effect. A Type III ANOVA was then conducted using the ‘car’ package.

## Results

### Age at reproductive onset

Treatment significantly impacted the age at reproductive onset (χ^2^_(2)_ = 42.50, *p* < 0.0001, Figure 1). Post-hoc comparisons showed that IG replicates started reproducing significantly earlier (mean ± SD: 80.1 ± 29.6 days) than both the I (104.6 ± 22.1 days) and T (109.5 ± 15.9 days) replicates (both *p* < 0.0001). By contrast, there was no difference in reproductive timing between the I and T replicates (*p* = 0.49).

**Figure 1.**
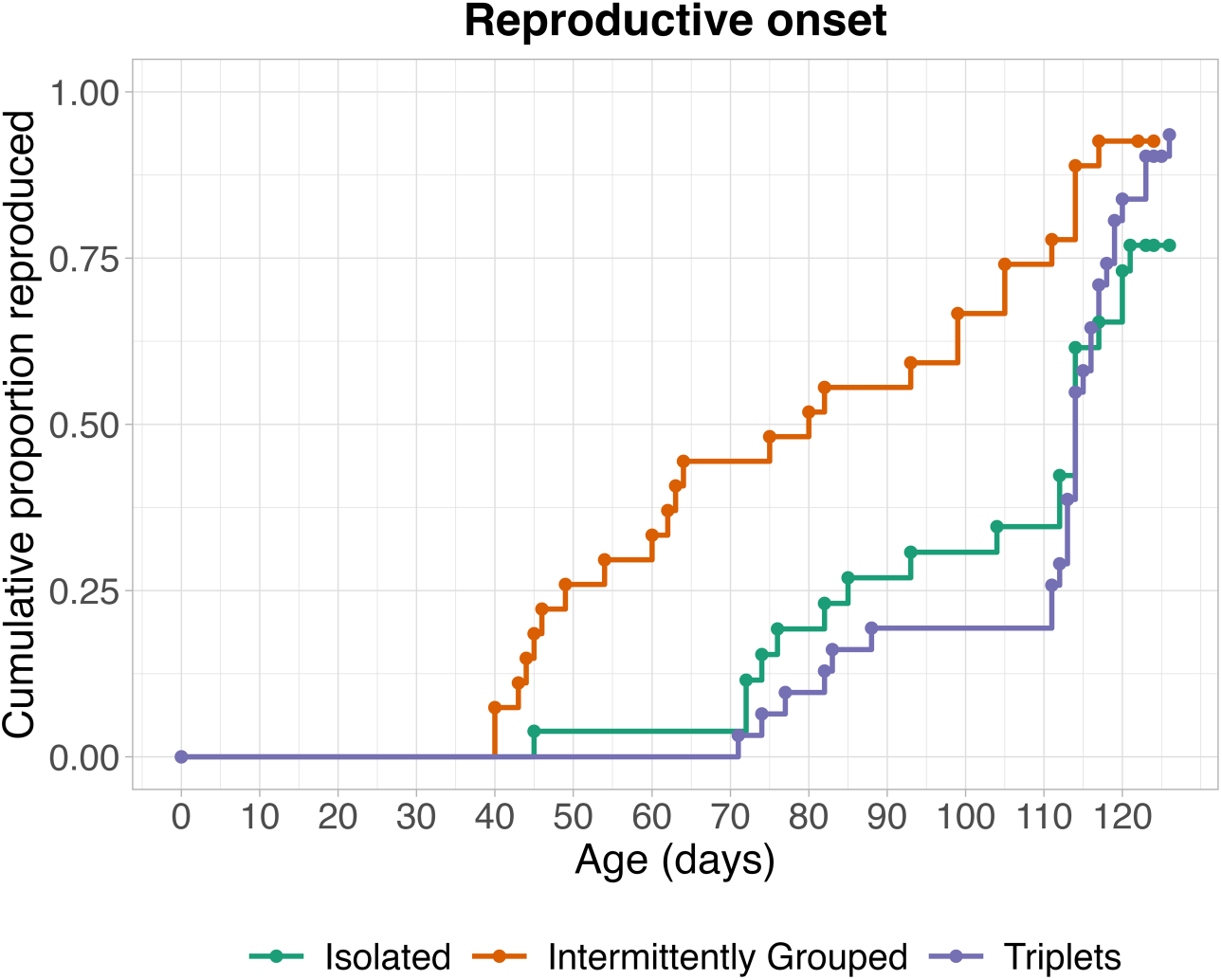
*Macrostomum hystrix* individuals with intermittent access to outcrossing opportunities (Intermittently Grouped) started reproducing significantly earlier (see *Results*) than worms with constant access to outcrossing opportunities (Triplets) or worms forced to self-fertilise (Isolated). The graph shows the cumulative proportion of replicates that began reproduction as a function of age for each treatment group. Mean ± S.D. values of age at reproductive onset (in days) for Isolated: 104.6 ± 22.1, Intermittently Grouped: 80.1 ± 29.6 and Triplets: 109.5 ± 15.9.

### Hatchling-to-adult survival

22 hatchlings were excluded from the analysis because they were younger than 21 days old when checked. Hatchlings were monitored from 16 I, 26 IG, and 24 T replicates, totalling 243 hatchlings. Treatment had no significant impact on hatchling-to-adult survival (*F*_2, 63_ = 0.1123, *p* = 0.89; Figure 2). Since offspring hatched over a long period of the experiment, we also compared survival for a restricted set of *n* = 148 offspring that all hatched within a seven day period when all three treatment groups were productive (days 121 – 127), with qualitatively identical results (*F*_*2,49*_ = 0.2264, *p* = 0.79; data not shown).

**Figure 2.**
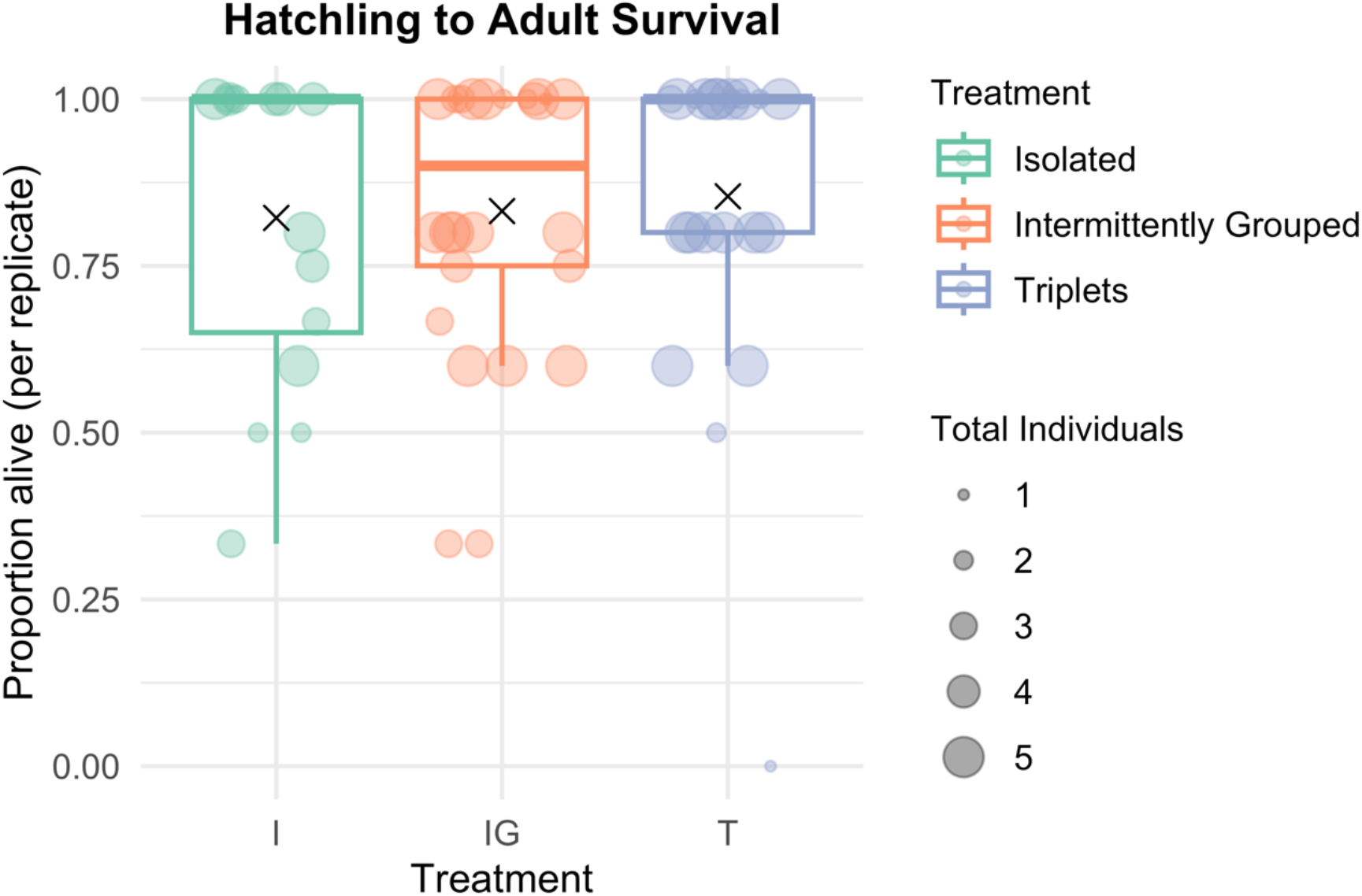
Progeny (F1) survival to adulthood did not differ between treatment groups (see *Results*). Up to 5 hatchlings were collected per replicate and each point shows the proportion that survived to adulthood, with point size reflecting variation in the number of F1 offspring monitored per F0 adult. Mean values are shown as black crosses. Sample sizes (F0 individuals) are *n* = 16 (Isolated), *n* = 26 (Intermittently Grouped) and *n =* 24 (Triplets).

## Discussion

Our findings suggest that a *Macrostomum hystrix* inbred line has a higher selfing propensity compared to its ancestral population. Following several generations of selfing, and despite an intervening period of relaxed selection, we observed a complete purging of inbreeding depression and the absence of a waiting time before selfing in this line. For mixed maters, whether selfing or outcrossing is the fertilisation mode employed within a generation is highly influenced by the environment [38]. Selfing, despite normally carrying fitness costs, is advantageous under conditions of low pollinator or mating partner availability, as it potentially allows preferentially outcrossing populations to tide over such periods and might partially explain the existence of mixed-mating strategies [2, 39]. However, if these conditions persist, repeated selfing is expected to lead to a shift in a population’s mating system [13, 19] and our results support this prediction.

The absence of inbreeding depression and waiting time in our experiment matches the theoretical predictions for this level of inbreeding [11]. These results are in line with experimentally evolving lines of *P. acuta* [19], although these authors found a reduction in inbreeding depression and waiting time rather than a complete elimination, likely because their inbreeding regime was less severe (but note also that recent evidence in *P. acuta* suggests that inbreeding depression can persist or be rapidly regenerated even in the absence of genetic variation [40]). More broadly, there is a tendency for mechanisms and behaviours to become more permissive towards future selfing in proportion to the amount of inbreeding depression purged. For example, 10 generations of selfing caused a greater reduction in anther-stigma distance compared to 10 generations of mixed mating in *Mimulus guttatus* [41]. Such rapid mating system shifts are particularly relevant for colonisations and expansions. Indeed, the ability of a single individual to found a new population is a hallmark characteristic of self-compatible species, and at least at some stage of the expansion or colonisation process, high selfing propensity is expected to be favoured by selection [16]. It is quite likely that our experimental regime created direct selection for early reproduction since we started with a single fastest-to-self founder and then selected a single individual each generation during the selfing phase. Although extreme, these conditions still fall within the range of possible scenarios that we might envisage in nature.

A second aim for our experiment was to partition the potentially negative social effects of constant grouping from differences arising from variation in mating system expression *per se*. We therefore quantified the consequences of constant versus intermittent grouping in *M. hystrix*. While we found no differences in progeny survival, individuals grouped intermittently started reproducing significantly earlier than those that were always grouped.

This could be interpreted as a lack of grouping costs in the IG treatment, although that alone wouldn’t explain why the I treatment (isolated) worms did not reproduce as early as the IG worms. One pertinent feature of the IG treatment compared to the other two is that at each grouping event, every individual’s housing well got shuffled and its group size changed for 2 hours every 2^nd^ or 3^rd^ day. Hence, one hypothesis to explain these differences is that this shuffling acted as a cue of an unstable social and/or physical environment in which future reproduction would not be guaranteed, thus triggering early reproduction. This early reproduction presumably comes at some costs otherwise we would expect all individuals in the population to reproduce sooner, but whether early reproduction trades off with future or total reproductive capacity is something that remains to be determined.

In conclusion, despite the fact we might have expected variation in reproductive onset to have been eroded by the selection regime, intermittent grouping of *M. hystrix* individuals nevertheless led to significantly earlier reproduction than either of the other two treatments.

Further studies should clarify whether this behaviour is primarily driven by the social or the physical component of the environment and more fully assess its fitness consequences, but our results establish that environmental effects on reproductive timing persist even after strong selection to eliminate variation in waiting times and inbreeding depression.

## Acknowledgements

This research was partly funded by the Agence Nationale de la Recherche (ANR), grant ANR-22-CPJ2-0116-01. A CC BY license is therefore applied to any AAM arising from this submission, in accordance with the grant’s open access conditions. We thank L. Schärer for maintaining a culture of the SR1 line over several years prior to this experiment.

